# A murine commensal protozoan relieves *Clostridium difficile* infection through regulating the arginine metabolism and the host intestinal immune response

**DOI:** 10.1101/2023.05.06.539719

**Authors:** Huan Yang, Xiaoxiao Wu, Wanqing Zang, Yuan Zhou, Xiao Li, Liang Wang, Wenwen Cui, Yanbo Kou, Yugang Wang, Bing Gu

## Abstract

Antibiotic-induced dysbiosis is a key predisposing factor for *Clostridium difficile* infection (CDI), fecal microbiota transplantation (FMT) is recommended for treating CDI. However, the mechanisms remain unclear. Here, we uncover that one integral member of the murine gut microbiota, a commensal protozoan from mice, *Tritrichomonas musculis* (*T. mu*), that reduced intestinal damage by inhibiting the recruitment of neutrophil and secretion of IL-1β, and increased Th1 cell differentiation and secretion of IFN-γ, which increased the goblet cell and secretion of mucin to protect the intestinal mucosa. The *T. mu* upregulated the key enzymes of arginine metabolism iNOS, ASS, OTC and down-regulated ARG1 in CDI mice, resulting in an increase in arginine, citrulline and a decrease in ornithine, which worked together to regulate the host intestinal immune response and alleviate CDI. This work reveals the interaction mechanism between intestinal commensal eukaryotes and pathogenic bacteria, and also provide new ideas for treating CDI.

## Introduction

*Clostridium difficile* (*C. difficile*) is a gram-positive anaerobic bacillus. *C. difficile* spore, conferring resistance to adverse environment and allowing persistence for several months, is the main pathogen of antibiotic-associated diarrhea^1^. *Clostridium difficile* infection (CDI) is widespread worldwide and imposes a huge economic burden on society^2^. Antibiotics are the main therapy for CDI, but due to overuse of antibiotics, *C. difficile* has developed a strong tolerance to a variety of antibiotics, including clindamycin, aminoglycosides and lincomycin^3^. Antibiotics such as metronidazole, vancomycin and fedamycin are still effective in treating CDI. But there are also some disadvantages, such as adverse reactions, recurrent CDI, and multi-resistant strains^4–6^. Therefore, more attention has been paid to non-antibiotic treatment of CDI in recent years.

Emerging non-antibiotic therapies mainly include fecal microbial transplantation (FMT)^7, 8^, probiotics^9–11^, diet^12–14^, etc. Different from the bactericidal effect of antibiotics, these therapies are more likely to regulate intestinal microflora to inhibit the germination and growth of *C. difficile* spores and enhance host immunity to reduce the damage caused by *C. difficile* ^7, 15, 16^. These emerging non-antibiotic therapies are more effective in treating recurrent CDI. FMT, widely noted, is a therapy that extracts and purifies microorganisms from the feces of healthy volunteers and then delivers them to the intestines of patients^7^. Most studies on the role and mechanism of FMT have focused on bacteria in feces, while the role of other microorganisms has not received enough attention.

The mammalian gut is home to a variety of microorganisms, including bacteria, fungi, protozoa, viruses, etc. Under normal circumstances, these gut microbes help host digestion and enhance host immunity against invasion of pathogen^17^. Most studies on the probiotic effects of intestinal microorganisms come from prokaryotes such as bacteria, while eukaryotes such as fungi, worms and protozoa are often found to have adverse effects on the host. *Entamoeba histolytica*^18, 19^ *Giardia lamblia*^20^ and *Cryptosporidium*^21, 22^ often cause disease with diarrhea as the main clinical manifestations. While the parasite may have unexpected protective effects. For example, *Cryptosporidium tyzzeri,* a cryptosporidium isolated in a laboratory at Washington University, can reside in healthy wild-type mice for a long time without causing symptoms, and enhance the colon Th1 response against *Citrobacter* infection^23^. In addition, in 2016, Chudnovskiy et al. discovered a commensal protozoan *Tritrichomonas musculis* (*T. mu*) in the healthy mice. This protozoan activates host intestinal epithelial inflammasome, induces IL-18 release, and promotes dendritic cell-driven Th1 and Th17 immunity^24, 25^.

At present, the mechanism of action of *T. mu* mainly focuses on *T. mu* immune regulation of host^24^. Existing studies have shown that in addition to intestinal immunity, the composition of intestinal microbes in *T. mu*-colonized mice has also changed^26^. Changes in the gut microbiome are often accompanied by changes in metabolism. At present, the research on the effect of *T. mu* on host metabolism only involves glucose metabolism^27^. Feces of patients infected with *C. difficile* contain rich 4-methylvaleric acid, the fermentation product of leucine, suggesting that *C. difficile* tends to use amino acids for energy in CDI^28^. Anaerobic protozoa derive energy mainly from carbohydrate and amino acid metabolism^29^. Studies have shown that the arginine dihydrolase pathway of anaerobic protozoa has the ability to provide ATP to cells^30, 31^. Therefore, there may be amino acid metabolism crosstalk between *T. mu* and *C. difficile*. Here, we introduced *T. mu* into the CDI model, aiming to explore the mechanism of intestinal commensal eukaryote *T. mu* on CDI, which is helpful to clarify the therapeutic principle of FMT and develop biologic agents for the treatment of CDI.

## Results

### 1. *T. mu* alleviates clinical manifestations and intestinal damage in CDI mice without affecting *C. difficile* colonization

Scanning electron microscopy showed that *T. mu* was pear-shaped with three anterior flagella and one posterior flagella (Supplementary Fig. 1A). First, to observe the effects of *T. mu* on healthy mice, mice were given an intragastric administration of purified *T. mu* every two days for a week, and their body weight was monitored. In addition, because we used protozoan-killing antibiotics such as metronidazole and clindamycin in the CDI model, we set up two additional antibiotic treatment groups. We found that the weight change of mice given *T. mu* was similar to that of normal mice (Supplementary Fig. 1B), and the mice did not show clinical manifestations such as diarrhea and hunchback. In addition, more *T. mu* was colonized in antibiotic-treated mice (Supplementary Fig. 1C). These data showed that *T. mu* did not affect the normal growth of mice and antibiotics may be beneficial to *T. mu* colonization.

**Fig. 1.**
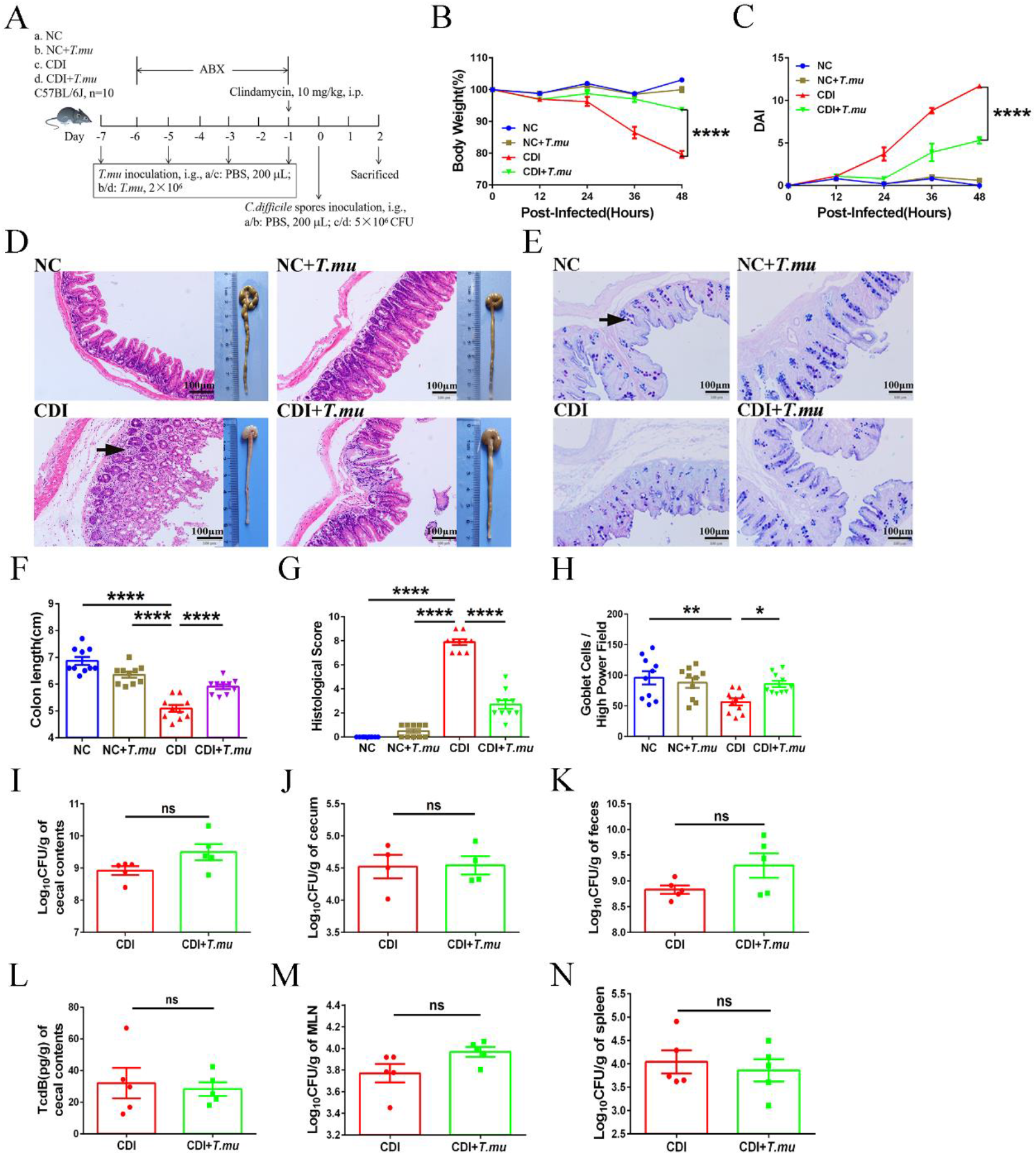
*T. mu* relieves CDI without affecting *C. difficile* colonization. **(A)** Schematic image illustrating co-colonization model of *T. mu* and *C. difficile*. *T. mu* negative WT mice were orally administered with purified *T. mu* every two days for a week, and then mice were infected with *C. difficile* spores. 48 hours later, cecum and colon of mice were collected. **(B)** Body weight loss. **(C)** Disease activity index (DAI). **(D)** Macroscopic findings of colon and representative HE-staining images (200×) of cecum. Scale bar: 100 μm. Arrow indicates infiltration of inflammatory cells. **(E)** Representative goblet cell staining images (200×) of cecum. Scale bar: 100 μm. Arrow indicates goblet cells. **(F)** Measurement and quantification of colon length. **(G)** Quantitation of histological score in cecum. **(H)** Quantification of cecal goblet cells. **(I)** Cecal contents, **(J)** cecal and **(K)** fecal *C. difficile* shedding. **(L)** The level of TcdB in cecal contents. The *C. difficile* load in **(M)** MLN and **(N)** spleen. Data are the mean ± SEM. n = 4-10. Statistical significance was determined by two-way ANOVA (**B** and **C**), one-way ANOVA (**F-H**) or Student’s t-test (**I-N**). ns no significance. *\**: *p<*0.05. *\*\**: *p<*0.01. *\*\*\**: *p<*0.001. *\*\*\*\**: *p<*0.0001.

Next, to observe the effect of *T. mu* on *Clostridium difficile* infection, we established a co-colonization model of *T. mu* and *C. difficile* (Fig. 1A). Within 48 hours after infection with *C. difficile*, we found that *T. mu* significantly alleviated weight loss, diarrhea, and humpback in CDI mice (Fig. 1B), and thus significantly reduced Disease activity index (DAI) (Fig. 1C). After 48 hours, the cecum and colon in CDI mice showed severe atrophy of cecum and colon, loss of intestinal epithelium, reduction of goblet cells, infiltration of inflammatory cells in intestinal mucosa leading to edema (Fig. 1D, E). In contrast, mice treated with *T. mu* had longer colons (Fig. 1F), lower histopathological score (Fig. 1G), and more goblet cells (Fig. 1H). These results suggest that *T. mu* can significantly relieve *C. difficile* infective colitis.

Based on the effect of *T. mu* on the CDI, we hypothesized that the *T. mu* may influence *C. difficile* colonization *in vivo*. Therefore, we measured the bacterial load in enteral and extrenteral tissues of CDI mice. Compared with *C. difficile* infected mice, cecal contents (Fig. 1I), cecum (Fig. 1J), and fecal contents (Fig. 1K) in *T. mu*-treated mice were not significantly reduced, and toxin contents of cecal contents were also high (Fig. 1L). These results suggests that *T. mu* does not inhibit *C. difficile* colonization in the intestine. In addition, Mesenteric lymph nodes (MLN) (Fig. 1M) and spleen (Fig. 1N) showed that *T. mu* did not affect the extrenteral spread of *C. difficile*. Thus, it is possible that effect of *T. mu* on CDI does not lie in inhibiting the colonization of *C. difficile*.

### 2. *T. mu* reduces neutrophil recruitment and cytokines secretion during CDI

It has been shown that *T. mu* affects the immune status of the host intestine^24^. Neutrophils have been shown to be critical early in the host response to *C. difficile*^32^. Therefore, we first focused on the infiltration of neutrophils in the colon, and the results were shown in Fig 2. *T. mu* did not promote neutrophil increase when colonized in mice for one week, while reduced the percentage of neutrophils in the gut after CDI (Fig. 2A). Correspondingly, we also found that *T. mu* significantly reduced the expression of neutrophil chemokine CXCL1 (Fig. 2B, C) and cytokines IL-36G, IL-1β and IL-6 (Fig. 2D-I), which may be the reason why *T. mu* alleviated intestinal inflammation in CDI mice. Next, we sought to determine whether *T. mu* reduced neutrophil production of IL-1β by *C. difficile*. *T. mu*, *C. difficile* and neutrophils were co-cultured *in vitro*, and the results showed that *T. mu* inhibited the secretion of IL-1β from neutrophils stimulated by *C. difficile* (Fig. 2J). These data suggest that *T. mu* may relieve *C. difficile* enteritis by reducing the recruitment of neutrophils and the secretion of pro-inflammatory cytokines in CDI mice.

**Fig. 2.**
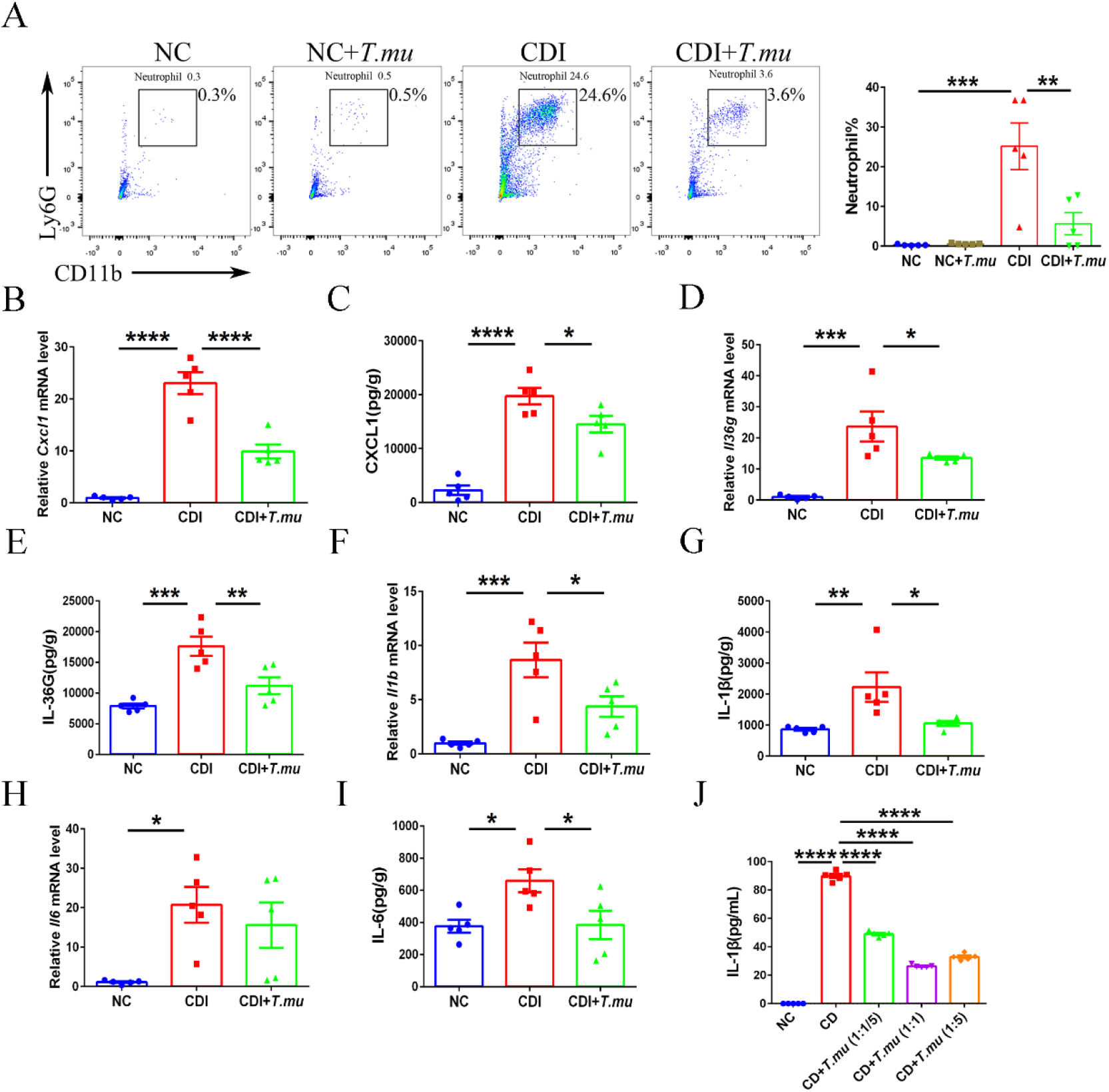
*T. mu* reduced intestinal neutrophil and pro-inflammatory cytokines in CDI. **(A)** Representative flow cytometry analysis of the neutrophils percentage in colonic lamina propria. **(B)** *Cxcl1*, **(D)** *Il36g*, **(F)** *Il1b* and **(H)** *Il6* levels were determined in the colon using qPCR. **(C)** CXCL1, **(E)** IL-36G, **(G)** IL-1β and **(I)** IL-6 levels were determined in the cecum using ELISA. (I) *In vitro*, different concentrations of *T. mu* (1:1/5, 1:1, and 1:5 *C. difficile*/*T. mu* ratios, respectively) reduced IL-1β of neutrophil stimulated by *C. difficile*. Data are the mean ± SEM. n = 5-6. Statistical significance was determined by one-way ANOVA. *\**: *p<*0.05. *\*\**: *p<*0.01. *\*\*\**: *p<*0.001. *\*\*\*\**: *p<*0.0001.

### 3. *T. mu* protects intestinal mucosa by increasing intestinal IFN-γ

Studies have shown that *T. mu* affects the differentiation of host colon T cells^24^. In our study, both CD4^+^IFN-γ^+^ (Th1) and CD8^+^IFN-γ^+^(Tc1) cells differentiated in the colons of healthy T. mu-colonized mice. After *C. difficile* infection, *T. mu* significantly increased the proportion of Th1 cells in the intestinal tract of mice (Fig. 3A, B). Consistent with the dominant Th1 cell induction, we observed a strong IFN-γ-driven gene and protein expression in the colon of *T. mu* colonized mice (Fig. 3C, D). These results prompted us to probe the effect of IFN-γ derived from Th1 cells in CDI. In addition, since parasites often enhance type II immunity in the host gut, we tested type II immune factors in *T. mu*-colonized mice here. The results showed that *T. mu* could not significantly increase the expression of intestinal cytokines IL-4 and IL-13 in colon (Fig. 3E, F). Therefore, the effect of *T. mu* on *C. difficile* infection may not be related to intestinal type II immunity.

**Fig. 3.**
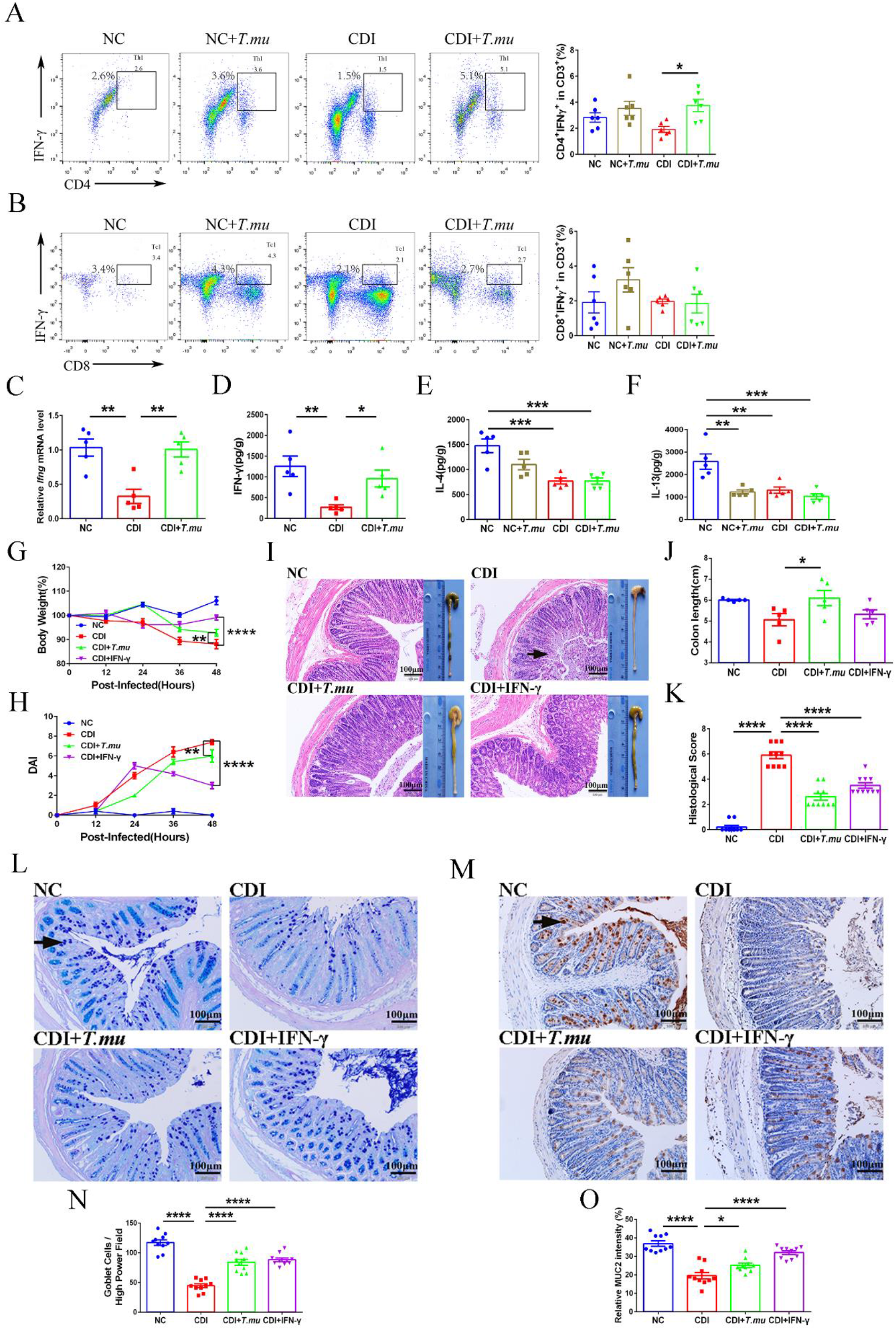
*T. mu* protects intestinal mucosa by increasing IFN-γ. Representative flow cytometry analysis of the **(A)** CD4^+^IFN-γ^+^ (Th1) and **(B)** CD8^+^IFN-γ^+^(Tc1) cells percentage in colonic lamina propria. **(C)** *Ifng* levels were determined in the colon using qPCR. **(D)** IFN-γ, **(E)** IL-4 and **(F)** IL-13 levels were determined in the cecum using ELISA. **(G)** Body weight loss. **(H)** DAI. **(I)** Macroscopic findings of colon and representative HE-staining images (200×) of colon. Scale bar: 100 μm. Arrow indicates infiltration of inflammatory cells. **(J)** Measurement and quantification of colon length. **(K)** Quantitation of histological score in colon. **(L)** Representative goblet cell staining images (200×) of colon. Scale bar: 100 μm. Arrow indicates goblet cells. **(M)** Representative MUC2 histochemical staining images (200×) of colon. Scale bar: 100 μm. Arrow indicates MUC2. **(N)** Quantification of colonic goblet cells. **(O)** Relative expression intensity of MUC2. Data are the mean ± SEM. n = 5-10. Statistical significance was determined by two-way ANOVA (**G** and **H**) or one-way ANOVA (**A-F**, **J**, **K**, **N** and **O**), *\**: *p<*0.05. *\*\**: *p<*0.01. *\*\*\**: *p<*0.001. *\*\*\*\**: *p<*0.0001.

Further, we injected the recombinant IFN-γ protein into CDI mice and found that IFN-γ improved their recovery from weight loss and diarrhea caused by *C. difficile* (Fig. 3G, H). Moreover, in IFN-γ-treated mice, the intestinal epithelium was relatively intact and histological scores were significantly reduced (Fig. 3I-K), which may be related to the enhanced obviously higher number of goblet cells with increased MUC2 protein level (Fig. 3L-O). These observations indicate that *T. mu* maintains the integrity of intestinal mucosa by increasing the expression of IFN-γ in the colon of CDI mice.

### 4. *T. mu* regulates arginine metabolism in CDI mice

Previous studies have shown that in CDI, *C. difficile* tends to use amino acids for energy^28^. Amino acid metabolism based on arginine dihydrolase pathway is one of the energy sources of anaerobic protozoa^30^, so there may be crosstalk of metabolism between *C. difficile* and *T. mu.* Performing KEGG pathways analysis, we found that intestinal metabolites of CDI mice mostly belong to amino acid metabolic pathways (Fig. 4A). Next, amino acids with differences between the two groups were enriched by KEGG pathway topological analysis, and the results showed that most of these amino acids with differences belonged to the arginine biosynthesis pathway (Fig. 4B). After correlation analysis of samples with different amino acids, the mice treated with *T. mu* showed higher intestinal abundance of citrulline and lower abundance of ornithine (Fig. 4C), and the two amino acids closely related to arginine biosynthesis (Fig. 4D). Measurement of amino acid concentrations in fecal samples revealed a significant increase of citrulline and arginine, while reduce the contents of ornithine in *T. mu* treated CDI mice compared with CDI mice (Fig. 4E-G). It may be related to the significant up-regulation of enzymes’ genes such as *iNos*, *Otc* and *Ass* (Fig. 4H-K), and the significant down-regulation of *Arg1* (Fig. 4L) in arginine metabolism of CDI mice treated with *T. mu*. In conclusion, the effect of *T. mu* on intestinal immunity in CDI mice may be related to the regulation of arginine metabolism.

**Fig. 4.**
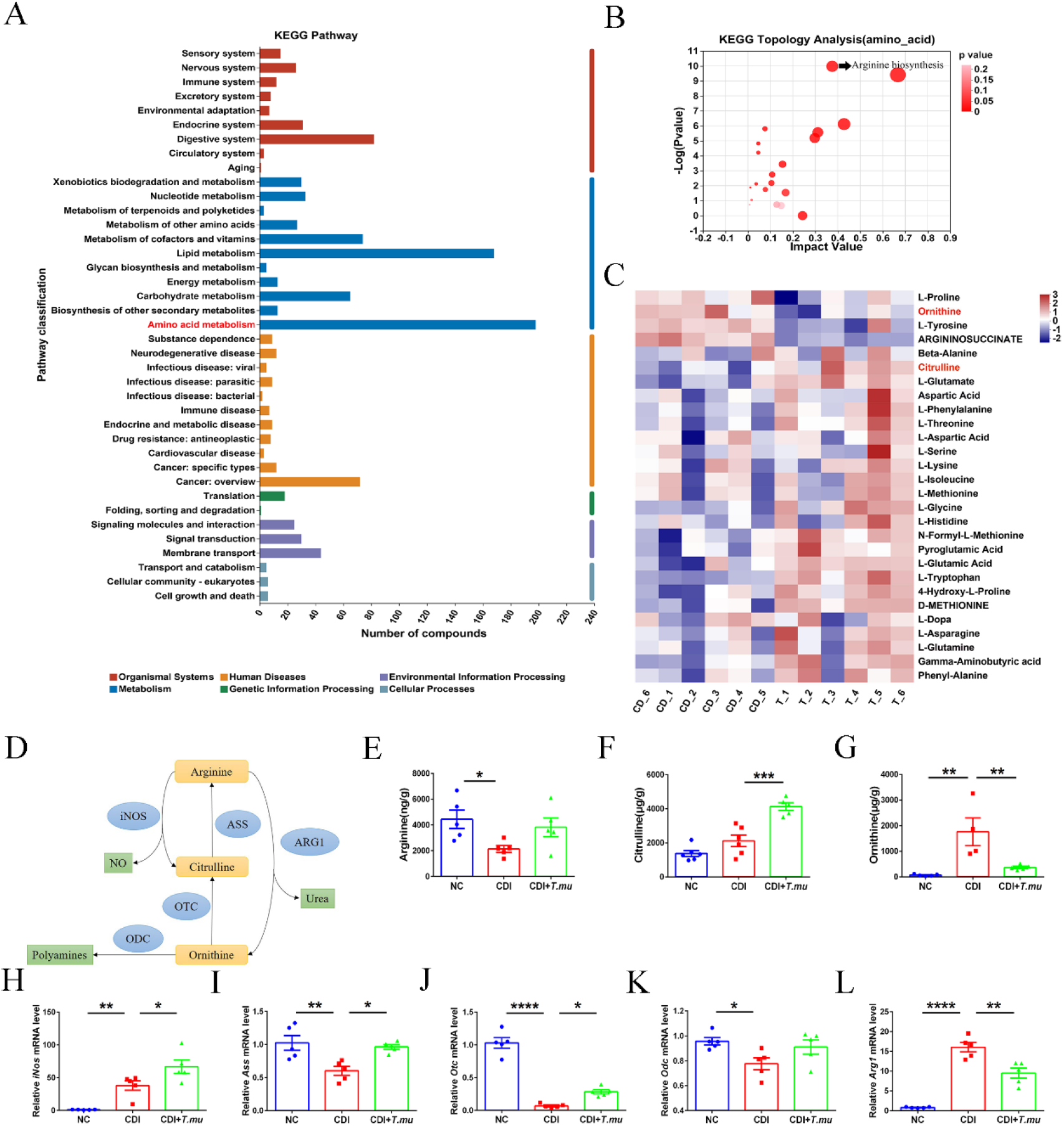
*T. mu* regulates intestinal arginine metabolism in CDI mice. **(A)** Statistical diagram of KEGG metabolic pathway. The ordinate is the secondary classification of KEGG metabolic pathway, and the abscissa is the number of metabolites annotated to this pathway. **(B)** KEGG pathway enrichment topology analysis. Each bubble represents a KEGG Pathway, the horizontal axis represents the impact value of the relative importance of metabolites in the pathway and the vertical axis represents the enrichment significance of metabolite participation pathway -log10(P value). **(C)** Correlation heat maps of Sample and amino acids with differences in the two groups. Each column in the Fig represents a sample, and each row represents a metabolite. The color in the Fig represents the relative expression quantity of metabolites in this group of samples. For the specific change trend of expression quantity, please see the digital label under the color bar on the upper right. **(D)** Diagram of arginine metabolism. Levels of **(E)** arginine, **(F)** citrulline and **(G)** ornithine in cecal contents. **(H)** *iNos*, **(I)** *Ass*, **(J)** *Otc*, **(K)** *Odc* and **(L)** *Arg1* levels were determined in the colon using qPCR. Data are the mean ± SEM. n = 4-6. Statistical significance was determined by one-way ANOVA (**E-L**). *\**: *p<*0.05. *\*\**: *p<*0.01. *\*\*\**: *p<*0.001. *\*\*\*\**: *p<*0.0001.

### 5. *T. mu* reduces neutrophil recruitment and IL-1β secretion by increasing intestinal arginine levels

In order to evaluate whether arginine metabolism is critical for *T. mu* to relieve CDI, the mice were fed citrulline and arginine in CDI mice. However, supplementation of citrulline did not reduce clinical symptoms or intestinal damage caused by *C. difficile* in mice (Supplementary Fig. 2A-E). Since citrulline can be converted to arginine under the regulation of ASS enzyme, we hypothesized that *T. mu* alleviated CDI through arginine. When supplemented with arginine, we found that arginine significantly reduced weight loss and diarrhea associated with CDI (Fig. 5A-B).

**Fig. 5.**
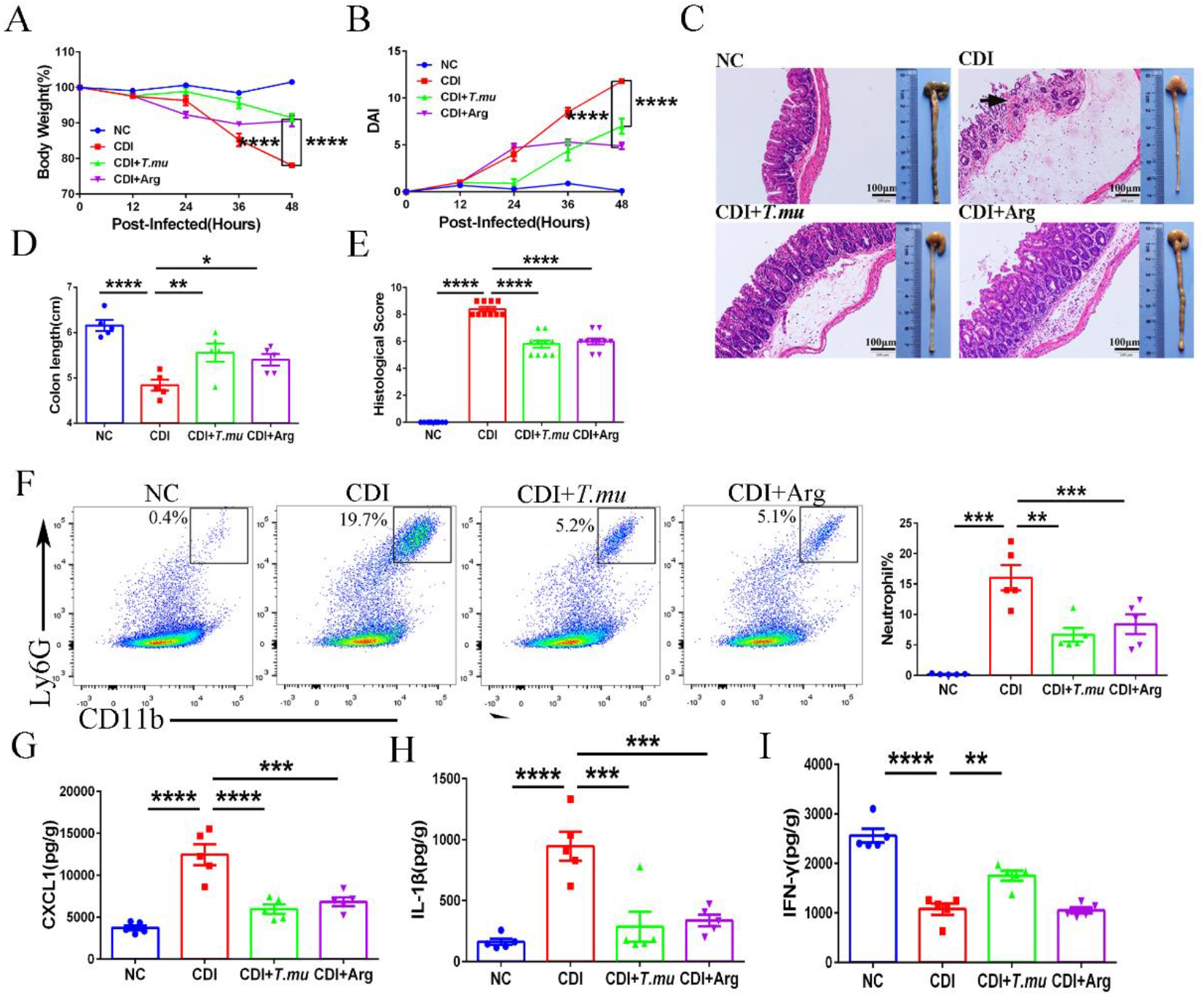
Arginine reduced the recruitment of intestinal neutrophils in CDI mice. **(A)** Body weight loss. **(B)** DAI. **(C)** Macroscopic findings of colon and representative HE-staining images (200×) of cecum. Scale bar: 100 μm. Arrow indicates infiltration of inflammatory cells. **(D)** Measurement and quantification of colon length. **(E)** Quantitation of histological score in cecum. **(F)** Representative flow cytometry analysis of the neutrophils percentage in colonic lamina propria. **(G)** CXCL1, **(H)** IL-1β and **(I)** IFN-γ levels were determined in the cecum using ELISA. Data are the mean ± SEM. Statistical significance was determined by two-way ANOVA (**A** and **B**) or one-way ANOVA (**D-I**). *\**: *p<*0.05. *\*\**: *p<*0.01. *\*\*\**: *p<*0.001. *\*\*\*\**: *p<*0.0001.

Additionally, in assessing intestinal injury in mice, we found that intestinal epithelial cell damage and inflammatory cell infiltration were reduced and cecal atrophy was alleviated in arginine-treated mice (Fig. 5C-E). Further, we found that arginine decreased the secretion of the chemokine CXCL1, the recruitment of neutrophils, and the expression of IL-1β in the intestinal tract of CDI mice. (Fig. 5F-H) However, it did not enhance IFN-γ-mediated immune protection in our study (Fig. 5I). These data collectively showed that arginine was associated with the relief of clinical symptoms of CDI and decreased recruitment of neutrophils in *T. mu* treated mice, while citrulline was not involved. Together, these results give strong support to the concept that protection in our model correlates with an increased availability of arginine.

### 6. *T. mu* reduces intestinal ornithine to regulate neutrophil recruitment and Th1 cell differentiation

As the levels of ornithine were significantly reduced in *T. mu*-treated mice, we hypothesize that ornithine may also play an important role in CDI. We further explored the effect of ornithine deficiency on CDI. Interestingly, in mice fed ornithine-free diet, there were almost no clinical symptoms of CDI (Fig. 6A, B), and intestinal damage was minimal (Fig. 6C-E). Then, we analyzed neutrophils and Th1 cells in the lamina propria of the colon. Compared with the CDI group, the proportion of neutrophils was significantly lower, while the proportion of Th1 cells was higher in the ornithine-free mice (Fig. 6F, G). And correspondingly, the expression of inflammatory factors CXCL1 and IL-1β was decreased, and the expression of IFN-γ was significantly increased after feeding ornithine-free diet for a week. (Fig. 6H-J). These data suggested that by reducing the ornithine content in the intestinal tract of mice, *T. mu* upregulated the expression of IFN-γ to protect intestinal mucosa from *C. difficile*, and thus reduces intestinal inflammation caused by neutrophils. These findings also demonstrate that modulation of metabolism and the immune response in the gut can alter CDI.

**Fig. 6.**
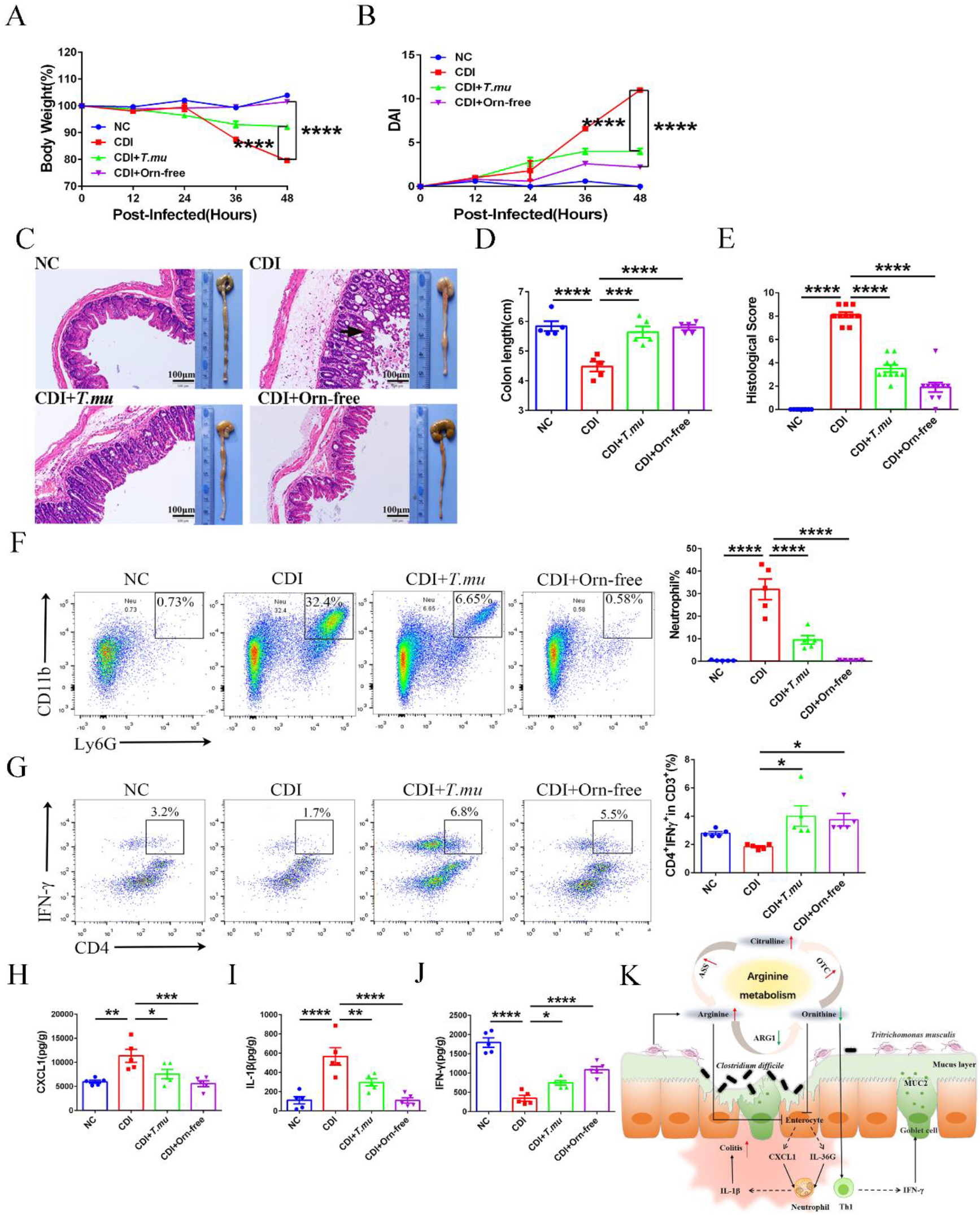
Ornithine-free diet decreased the recruitment of intestinal neutrophils and increased Th1 cell differentiation in CDI mice. **(A)** Body weight loss. **(B)** DAI. **(C)** Macroscopic findings of colon and representative HE-staining images (200×) of cecum. Scale bar: 100 μm. Arrows indicate infiltration of inflammatory cells. **(D)** Measurement and quantification of colon length. **(E)** Quantitation of histological score in cecum. Representative flow cytometry analysis of the **(F)** neutrophils and **(G)**Th1 percentage in colonic lamina propria. **(H)** CXCL1, **(I)** IL-1β and **(J)** IFN-γ levels were determined in the cecum using ELISA. **(K)** The hypothetical diagram of the mechanism of *T. mu* in treating CDI. Data are the mean ± SEM. n = 5-10. Statistical significance was determined by two-way ANOVA (**A** and **B**) or one-way ANOVA (**D-J**), *\**: *p<*0.05. *\*\**: *p<*0.01. *\*\*\**: *p<*0.001. *\*\*\*\**: *p<*0.0001.

## Discussion

In this study, protozoan *T. mu* was extracted from the cecum of *HVEM ^-/-^*mice. In addition to *HVEM ^-/-^* mice, *T. mu* has been detected in several commercial mice^24, 26^. However, as a parasite, *T. mu* did not cause pathological damage to the intestinal tract of mice, nor did it show clinical symptoms such as diarrhea, that is, *T. mu* showed symbiotic status in these mice. The antibiotics used in the CDI model did not kill *T. mu*, but helped to *T. mu* colonize. In order to explore the interaction mechanism between intestinal eukaryote *T. mu* and prokaryote *C. difficile*, we introduced *T. mu* into CDI mouse. The results showed that *T. mu* could effectively relieve intestinal injury in CDI mice, which was independent of reducing intestinal colonization of *C. difficile*. *C. difficile* toxins TcdA and TcdB damage the cytoskeleton of intestinal epithelial cells and increase intestinal permeability, trigger the release of chemokines CXCL1 and CXCL2, a large number of neutrophils will be activated at the site of infection, and produce more chemokines and cytokines. Which results in a strong neutrophil-mediated inflammatory response^33–35^. IL-36G belongs to the IL-1 family and can be hydrolyzed and activated by neutrophil protease, playing a key role in the pathogenesis of psoriatic skin inflammation^36, 37^. In inflammatory bowel disease, IL-36G was found to be expressed in intestinal epithelial cells, where the IL-36R signaling pathway control recruitment of white blood cells to the inflammatory site^38^. In *Candida albicans* infection^39^, Streptococcus pneumoniae^40, 41^, and herpes simplex virus type 2 ^42^, IL-36G promotes the recruitment of neutrophils to inflammatory centers and accelerates phagocytosis and clearance of pathogens. Our results showed that CDI also caused the high expression of IL-36G, which may be related to the recruitment of neutrophil-scavenging bacteria by IL-36G, and the inflammatory response caused by excess neutrophils aggravated the intestinal injury. Our work showed that *T. mu* can reduce intestinal inflammation through inhibiting the expression of cytokines IL-1β, IL-6 and IL-36G and the over recruitment of neutrophils.

It is known that *T. mu* colonization in mice significantly amplified Th1 and Th17 cells in colonic mucosa, but not Th2 cells ^24^. We uncover that *T. mu* can restore the Th1 cells counts and IFN-γ in the gut of CDI mice, while the type II immune cytokines, IL-4 and IL-13, were not significantly increased. Thus, *T. mu* -activated Th1 cells may help alleviate *C. difficile* infection. The role of IFN-γ in intestinal inflammation is controversial. IFN-γ has been previously reported to be involved in various types of chronic inflammatory diseases and TcdA-mediated acute enteritis ^43^. However, recent studies have found that IFN-γ produced by type I ILCs can restrict the replication of *Toxoplasma gondii* in the intestinal tract^44^ and prevent the invasion of *Clostridium difficile*^45^. Moreover IFN-γ secreted by Th1 cells has protective effects against *Cryptosporidium*^46^, Enterohemorrhagic *Escherichia coli*^47^, *Salmonella* ^48, 49^ and *Clostridium difficile*^50^. The IFN-γ-mediated antibacterial mechanism mainly includes stimulating Paneth cells to release antimicrobial peptides^51^. and regulating mucin production to protect intestinal mucosa^52^ and limit bacterial systemic transmission^53^. We find that MUC2 secreted by goblet cells significantly decreased in CDI and intestinal mucus with MUC2 as the main component mainly provided a physical barrier for intestinal epithelium, protecting it from pathogen infestation. Moreover, *T. mu* can increase Th1 cells and he expression of IFN-γ in CDI mice, and then up-regulate the secretion of MUC2, which protects intestinal mucosa.

Amino acid metabolism plays an important role in *T. mu*, and the arginine dihydrolase pathway can also provide energy for protozoa growth^29, 54^. *Clostridium difficile* also appears to be more dependent on amino acid metabolism for energy in CDI patients^28^. Our data show that *T. mu* significantly altered amino acid metabolism in CDI mice. Among them, KEGG pathway enrichment analysis reveals that arginine anabolic enrichment is the most significant. *In vitro* culture of protozoa can produce citrulline, which is further involved in arginine metabolism. *T. mu* upregulates *Ass* to convert citrulline to arginine, and down-regulated *Arg1* to inhibit the conversion of arginine to ornithine. In conclusion, the citrulline and arginine are increased in the intestinal tract of mice treated with *T. mu* in CDI, while the ornithine is decreased. Supplementation of citrulline cannot relieve the clinical symptoms of CDI mice, while supplementation of arginine can inhibit the recruitment of neutrophils and reduce the secretion of IL-1β, thus reducing intestinal inflammation. *T. mu* intervention reduces ornithine in *C. difficile* infected mice. Therefore, when given ornithine-free diet in CDI mice, Th1 cells in mice were significantly expanded, promoting IFN-γ secretion and maintaining the integrity of intestinal epithelium. And neutrophils recruitment and IL-1β secretion were rare in mice feed with ornithine-free diet to relieve inflammatory response. In summary, *T. mu* can alleviate CDI by affecting host immunity through regulating arginine and ornithine in arginine metabolism.

In conclusion, this study focused on intestinal metabolism and immunity to explore the mechanism of commensal protozoan *T. mu* on intestinal inflammation induced by *C. difficile*. Many studies have reported the harm of pathogenic *Amoeba* and *Flagellates*, while the probiotic effects of some symbiotic eukaryotes have been less mentioned. We propose a model of the interaction between commensal protozoan and *C. difficile*, in which *T. mu* regulates the host immune response by regulating arginine metabolism to maintain intestinal homeostasis and relieve *C. difficile* infection (Fig. 6K). Based on our data, we propose that *T. mu* intervention in the induced increase of arginine and decrease of ornithine in CDI mice leads to reduce intestinal damage and increase goblet cell secretion of MUC2 protein, in which neutrophils and Th1 cells play a critical role. In future studies, it may be necessary to detect the colonization of intestinal *T. mu* in healthy humans and CDI patients, and focusing on the role of eukaryotes in feces from healthy donors may help FMT therapy break through the bottleneck of ethical and safety threats.

## Methods

### Animals and Intervention

All mouse studies were evaluated by the Laboratory Animal Ethics Committee of Xuzhou Medical University (IACUC number: 202202A278, Xuzhou, China). Wild-type C57BL/6J mice (4 weeks old; male) were purchased from Xuzhou Medical University. The mice were housed on a 12 h alternating day and night cycle, and standard laboratory sterilized feed and water were freely available. The room temperature was 25°C, and the humidity was 40–70%. Groups of age and gender matched C57BL/6 mice were either left untreated or were orally gavaged with 2×10^6^ /mouse purified *T. mu*. For experiments, the 40 mice were randomly divided into four groups: normal mice without *T. mu* (designed NC group), normal mice treat with *T. mu* (designed NC +*T. mu* group). The other two groups exposure to *C. difficile*. One group were fed with *T. mu* (designed CDI +*T. mu* group), One group were fed with PBS (designed CDI group).

In the IFN-γ recombinant protein intervention experiment, each mouse in the IFN-γ intervention group (CDI+IFN-γ group) was intraperitoneally injected with 10 μg IFN-γ recombinant protein (SinoBiological, China) at 2 h and 24 h after *C. difficile* infection. In the arginine intervention experiment, mice in the arginine intervention group (CDI +Arg group) were intraperitoneally injected with 200 mg/kg arginine (Sangon, China) for 7 days before infection. In the citrulline intervention experiment, mice in the citrulline intervention group (CDI +Cit group) were given 2% citrulline (Sangon, China) solution for 7 days before infection. In the ornithine-free diet intervention experiment, mice in the ornithine deficiency intervention group (CDI +Orn-free group) were given ornithine-free diet (Xietong, China) for 7 days before infection. This diet formula refers to the published literature.^55^

### Purification of *T. mu*

The cecal contents of *T. mu* containing mice were harvested into sterile PBS and filtered several times through a 70 µm cell strainer. The suspension was spun at 1500 rpm for 5 min at 4 ℃. The pellet was washed twice with PBS. *T. mu* was further purified at the interface of a 40%/80% percoll (cytiva, USA) gradient. For *in vivo* administration, each mouse was inoculated with approximately 2×10^6^ T. mu via the oral route.

### Scanning Electron Microscopy

As previously^26^, purified *T. mu* was suspended and fixed with fixative (2.5% glutaraldehyde + 4% paraformaldehyde) in PBS for 2 h and washed three times. Then samples were dehydrated using the following series ethanol-water mixtures: 25%, 50%, 70%, 80%, 90%, 100%, 100%. After treatment with 100% ethanol the samples were incubated in isoamyl acetate twice. The samples were dried in a critical-point dryer HCP-2 (Hitachi Ltd., Japan). Then the samples were attached to the sample table and gold coating was performed in a high vacuum coating instrument (Leica EM ACE600, Germany). After gold coating, the samples were observed using a scanning electron microscope Teneo VS (FEI, USA)

### *C. difficile* Spore Preparation

*C. difficile* VPI 10463 (ATCC 43255) spores were prepared as described previously^56^. *C. difficile* VPI 10463 strain was cultivated for 2 days in Brain Heart Infusion broth (OXOID, England) supplemented with 1.5% agar and 0.05% L-cysteine at 37°C in anaerobic atmosphere (AnaeroPack, Japan) in jars. Then, *C. difficile* was grown in a 5-ml culture of liquid medium (BHI supplemented with 0.05% L-cysteine) at 37 ℃ anaerobically. 2 days later, the inoculum was added to 125 mL culture of liquid medium. The culture was incubated at 37℃ for 5 days anaerobically. Spores were harvested by centrifugation and washed with cold PBS at least three times. *C. difficile* spores were heat treated for 10 min at 70℃ to remove vegetative cell debris. Spore stocks were stored at -20℃ in sterile PBS. The agar medium to recover viable spores was BHIYT prepared as follows: 3.7% BHI (OXOID, USA), 0.5% yeast extract (Y, Sangon, China), 0.1% L-cysteine (Sigma, USA), 0.1% sodium taurocholate (T, Sangon, China), and 1.5% agar (Solarbio, China). This medium was autoclaved for 20 min at 121℃. The numbers of viable spores were recorded as CFU/mL.

### Model of *Clostridium difficile* infection

Mice infections were performed as follows. Mice were pretreated with antibiotic mixture (ABX, 0.4 mg/mL kanamycin, 0.035 mg/mL gentamicin, 0.035 mg/mL colistin, 0.215 mg/mL metronidazole, and 0.045 mg/mL vancomycin; Sangon, Shanghai, China) added to drinking water for 5 days. Next, mice received clindamycin (10 mg/kg, i.p.; Sangon, China). After 1 day, mice were infected with 5×10^6^ CFU of *C. difficile* spores by gavage. Mice were weighed and monitored twice daily Symptoms of disease (stool characteristics, weight loss, and decreased response to stimuli) were recorded and mortality was tracked. During the entire protocol with a disease activity index (DAI) that varied from 0 (normal) to 12, as described^57^. Briefly, DAI is based on clinical symptoms of stool characteristics, behavioral change, and percent weight loss. Each category is scored from 0 to 4, and the individual values are added to provide an overall score.

### Quantification of *T. mu* in the cecal content

The cecum of mice was cut longitudinally and weighed. The harvested cecal content was suspended in PBS (10 µL/mg). The protist was counted using a hemocytometer as previously reported. The total number and concentration of *T. mu* were calculated accordingly.

### Gross and histological assessment

The whole colon from the end of the cecum to the anus was photographed and the length of the colon was measured. For histological analysis, cecum was fixed in 4% paraformaldehyde and embedded into paraffin. Tissue sections (4 μm) were stained with hematoxylin and eosin. Images were captured with a microscope (DP74, Olympus, Japan). Histological score was evaluated according to the criteria described previously^57^. Briefly, Histologic injury was graded based on epithelial tissue damage, amount of edema, and neutrophil infiltration. Each category was scored from 0 to 3 with the individual values added for an overall score.

### Enzyme-linked immunosorbent assay (ELISA)

Cecum samples were homogenized in RIPA Lysis Buffer (KeyGEN, China) containing protease inhibitors (Cwbio, China). Samples were centrifuged for 10 min at 10000 ×g, and supernatants used for measurement of IL-1β, IL-6, IL-13, IFN-γ, IL-4, IL-36G and CXCL1. IL-1β, IL-13, IFN-γ IL-4 and IL-6 were measured by Mouse Uncoated ELISA Kit (Invitrogen, USA) according to the manufacturer’s instructions. IL-36G were quantified using Mouse ELISA kits (DLDEVELOP, China) according to the manufacturer’s instructions. CXCL1 was quantified using Mouse ELISA kits (Fine, China) according to the manufacturer’s instructions.

### Real-time quantitative PCR (qPCR)

RNA from colonic tissues was extracted using the total RNA extraction kit (Solarbio, China) in accordance with the manufacturer’s instructions. One microgram of RNA was transcribed into cDNA using the PrimerScript RT Reagent Kit (Takara, Japan). After reverse transcription, qPCR amplification was performed using SYBR Green qPCR Master mixes (Bimake, USA) under thermocycler PCR system (LightCyler 96, Roche, Germany). The PCR program was as follows: 95 °C for 10 min; 40 cycles of 95 °C for 10 s, 60 °C for 30 s, and 72 °C for 32 s. 2^−ΔΔCT^method was used to calculate the relative gene expression. β-actin was used as an internal control. The primers used in this experiment were designed by Sangon Biotech Co., Ltd., (Beijing, China). The primers are listed in the Table 1.

**Table. 1.**
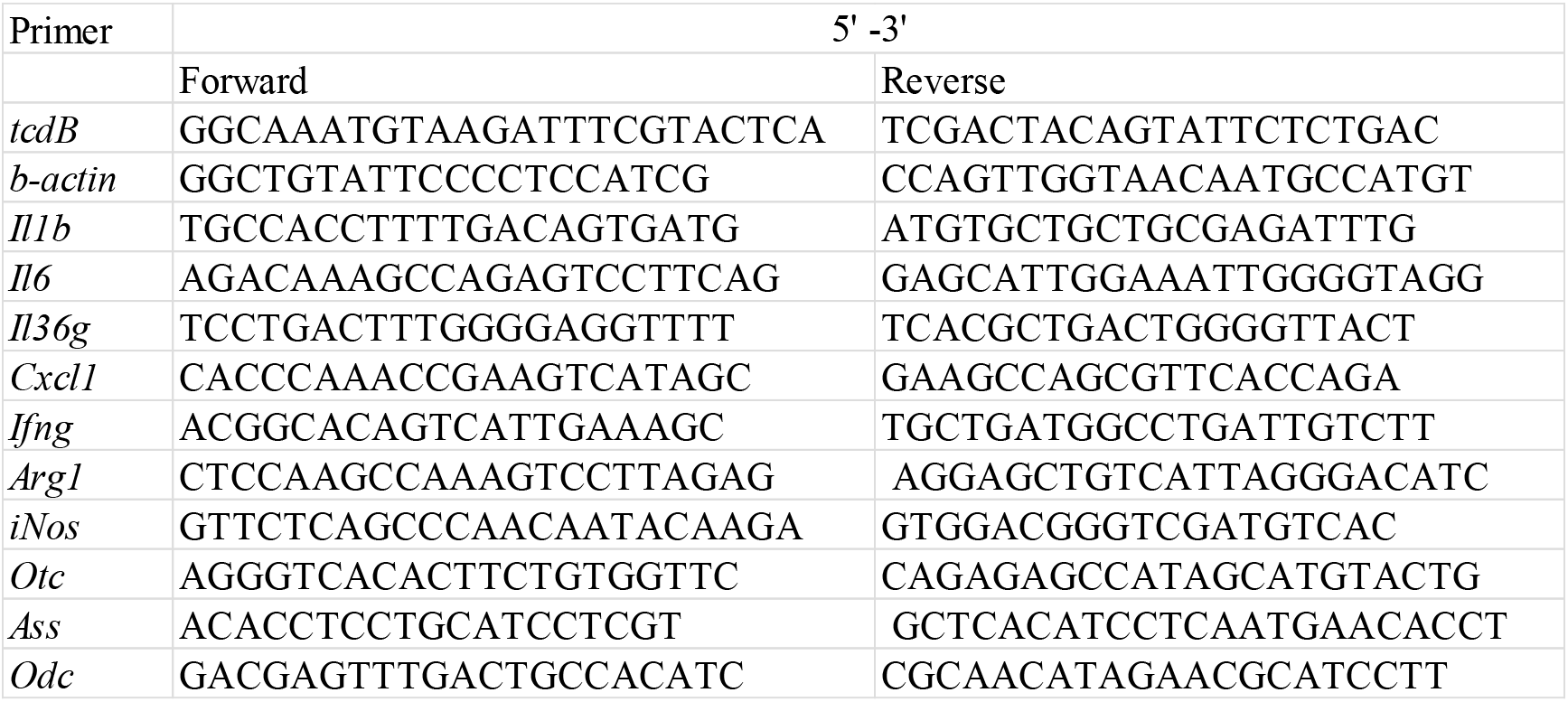
Primer sequences used in this study.

### Quantification of *C. difficile* and toxins

Levels of C. difficile toxins B in cecal contents and feces were measured by ELISA using the Mouse CDT-B ELISA KIT (Mlbio, China). For cecum, mesenteric lymph nodes, and spleen, the tissues were weighed and homogenized in 1mL of 0.3%Triton-100 solution, followed by 10-fold serial dilutions plated on BHI agar, and CFU per g of tissue was counted. For cecal contents and feces, fecal bacterial genomic DNA extraction kit (TIANGEN, China) was used to extract fecal *C. difficile* DNA after weighing, and then qPCR was used for quantification. *C. difficile* genomic DNA were extracted using the TIANamp Bacteria DNA Kit (TIANGEN, China), and gene levels were quantified by qPCR using TcdB primers (Table 1). Copy numbers of unknown sample were extrapolated from the standard curve that was generated with the extracted DNA prepared from known *C. difficile* inocula.

### Tissue preparation and flow cytometry

To isolate lymphocytes from the colonic lamina propria, the colon tissues were cut into 1-cm pieces in cold PBS and incubated with cold PBS containing 2 mM dithiothreitol (DTT) at 240 rpm and 37 °C for 10 min, followed by incubating with cold PBS containing 5 mM EDTA three times at 240 rpm and 37 °C for 10 min. Next, the colon tissue was minced with scissors in 1 mL digestion solution (1 mg/mL collagenase (Sigma-Aldrich, USA), 1 mg/mL hyaluronic acid (Sigma-Aldrich, USA), and 1μg/mL DNase I (MACKLIN, China) in PBS at 100 rpm and 37 °C for 40 min. After incubation, the cell solution was strained through a 70-μm cell strainer and suspended in 40% Percoll (cytiva, USA), followed by centrifugation at 670×g for 30 min at 4 °C. The cell pellet was washed with cold PBS and resuspended in 2% fetal bovine serum (ExCell, Australia). For individual surface antibody staining, single-cell suspensions were stained for 40 min at 4 °C in the dark with the surface antibodies. For intracellular staining, percoll isolated cells were activated for 5 hours at 37 °C using a Cell Activation Cocktail (1:500, Biolegend, USA). The cells were washed with PBS once and stained with surface staining antibody for 40min at 4 °C. The cells were washed with PBS twice, 1500rpm for 5min each time. Cells were immobilized with cell fixation buffer for 30min at 4 °C. The permeabilization buffer (Biolegend, USA) was used to permeate the cells twice, 1500rpm for 5min each time. Next, they were stained with intracellular staining antibodies for 5 hours at 4 °C. Cells were washed with permeabilization buffer and PBS successively. The cells were fixed with 0.2% paraformaldehyde and analyzed by flow cytometry. Surface staining antibodies: anti-CD45 conjugated to PE/Cyanine7 (1:200, Biolegend, USA), anti-CD4 conjugated to FITC (1:200, Biolegend, USA), anti-CD11b conjugated to Pacific Blue (1:200, Biolegend, USA), anti-Ly6G conjugated to PE (1:200, Biolegend, USA), anti-CD3 conjugated to PerCP/Cyanine5.5 (1:200, Biolegend, USA), anti-CD8 conjugated to APC (1:200, Biolegend, USA). Intracellular staining antibodies: anti-IFN-γ conjugated to Brilliant Violet 510 (1:200, Biolegend, USA), anti-IL-17A conjugated to Brilliant Violet 421 (1:200, Biolegend, USA).

### *In vitro* co-culture

Neutrophils were recovered from bone marrow suspensions following double-gradient centrifugation (Histopaque-1119, Histopaque-1077: 50/50 v/v; SIGMA, USA). Cells (1 × 10^5^ cells/mL) were plated at 37°C in DMEM (VICMED, China) containing 10% fetal bovine serum (ExCell Bio, Uruguay) and without antibiotics. Initially, cells were incubated for 2 h with 2 × 10^6^, 1 × 10^7^ or 5 × 10^7^ *T. mu*. *C. difficile* (1 × 10^7^ CFU/mL) was then added to the culture for 4 h. Subsequently, IL-1β cytokine released in the culture was quantified by ELISA.

### Quantification of Amino Acids

0.1g of cecal contents were suspended in 1ml of extraction solvent containing 0.1% formic acid (acetonitrile: methanol: water=2:1:1). After shaking, homogenizing and ultrasound, the suspension was incubated at – 20 ℃ for 1h. After centrifugation at 12000 × g for 15 min, supernatants were used for LC/MS.

### Untargeted metabolomics analyses

Liquid chromatography-mass spectrometry (LC-MS)-based metabolomics was performed by Majorbio Biotech (Shanghai, China). Briefly, a 50 mg cecal contents sample was accurately weighed, and the metabolites were extracted using 400 µL of methanol: water (4:1, v/v) solution. The mixture was allowed to settle at −10 ℃ and treated with a high-throughput tissue crusher Wonbio-96c (Shanghai Wanbo Biotechnology Co., Ltd, Shanghai, China) at 50 Hz for 6 min, followed by vortex for 30 s and ultrasound at 40 kHz for 30 min at 5 ◦C. The samples were placed at −20 ℃ for 30 min for precipitating the proteins. After centrifugation at 13000 × g and 4 ℃ for 15 min, the supernatant was carefully transferred to sample vials for LC-MS analysis. Chromatographic separation of the metabolites was performed using a Thermo UHPLC system equipped with an ACQUITY UPLC HSS T3 column (100 mm × 2.1 mm i.d., 1.8 µm; Waters, Milford, USA). Mass spectrometric data were collected using a Thermo UHPLC-Q Exactive HF-X Mass Spectrometer equipped with an electrospray ionization source operating in either a positive or negative ion mode. The data were analyzed through the free online platform of Majorbio cloud platform (cloud.majorbio.com). Metabolic features detected at least 80 % in any set of samples were retained. After filtering, minimum metabolite values were imputed for specific samples in which the metabolite levels fell below the lower limit of quantitation and each Metabolic features were normalized by sum. In order to reduce the errors caused by sample preparation and instrument instability, the response intensity of the sample mass spectrum peaks was normalized by the sum normalization method, and the normalized data matrix was obtained. At the same time, variables with relative standard deviation (RSD) > 30% of QC samples were removed, and log10 logarithmization was performed to obtain the final data matrix for subsequent analysis.

### Statistical analysis

Statistical analyses were performed using GraphPad Prism version 6.0 (GraphPad software, San Diego, CA). Data was presented as mean ± standard error of the mean (SEM). Significance in all cases was set at *p<*0.05. Scores, weights, and assay values were analyzed using a One-way or Two-way Repeated Measures Analysis of Variance (ANOVA) test, student’s two-tailed where appropriate. With bacterial quantification assays, the log10 values of the results were used to achieve normal distribution.

## Author Approvals

All authors have seen and approved the manuscript, which has not been accepted for publication elsewhere.

## Data availability statement

All data generated in this study are included in this article and supplementary information or are available from the corresponding author on reasonable request.

## Disclosure statement

The authors declare no competing interests.

## Author contributions

H.Y., X.W., Y.K. and Y.W. designed the study and prepared the manuscript. X.W., X.L., W.Z. and

H.Y. performed and analyzed experiments. X.W., H.Y., X.L., L.W., Y.Z. and Y.K. contributed intellectually to the analysis and interpretation of the data. X.W., W.C. and H.Y. wrote the manuscript. H.Y., B.G. and Y.W. provided funding for the research.

## Funding

This research was supported by the National Natural Science Foundation of China (82102408, 81871734, 82072380), China Postdoctoral Science Foundation(2022M712681), Research foundation for advanced talents of Guangdong Provincial People’s Hospital (KJ012021097) and Xuzhou Medical University Excellent Talent Introduction Project (D2019030, RC20552061).

---

**Supplementary Fig. 1.**
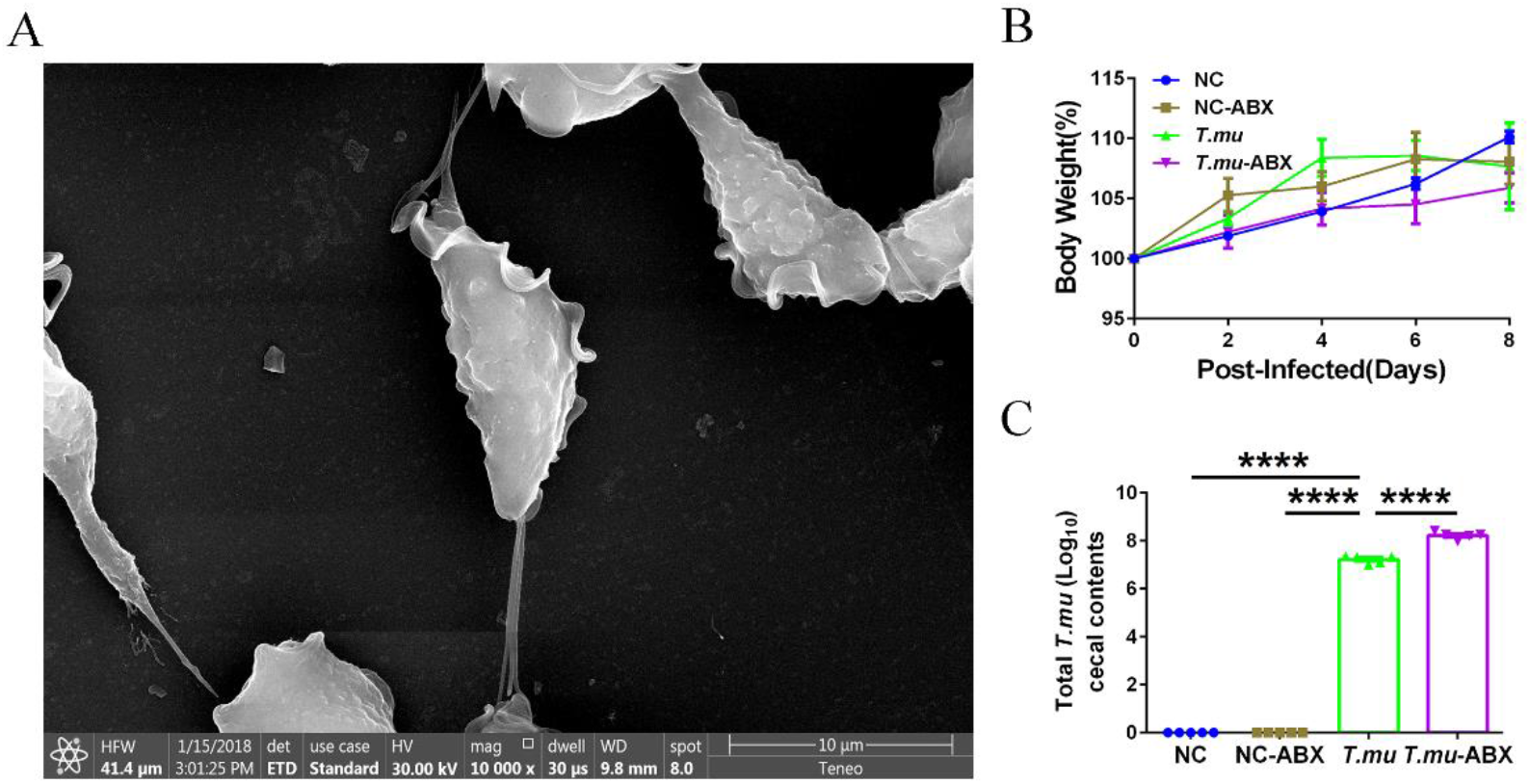
*T. mu* have no influence on the growth of mice. *T. mu* negative WT mice were orally administered with purified *T. mu* every two days for a week, along with an antibiotic mixture (ABX) for 5 days. At day 8, the mice were sacrificed to collect cecal contents. **(A)** Representative SEM image of purified *T. mu*, scale bar :10 μm. **(B)** Body weight changes in the indicated groups. **(C)** The total number of *T. mu* in the indicated groups. Data are the mean ± SEM. n = 5. Statistical significance was determined by two-way ANOVA (B) or one-way ANOVA (C). *\*\*\*\**: *p<*0.0001.

**Supplementary Fig. 2.**
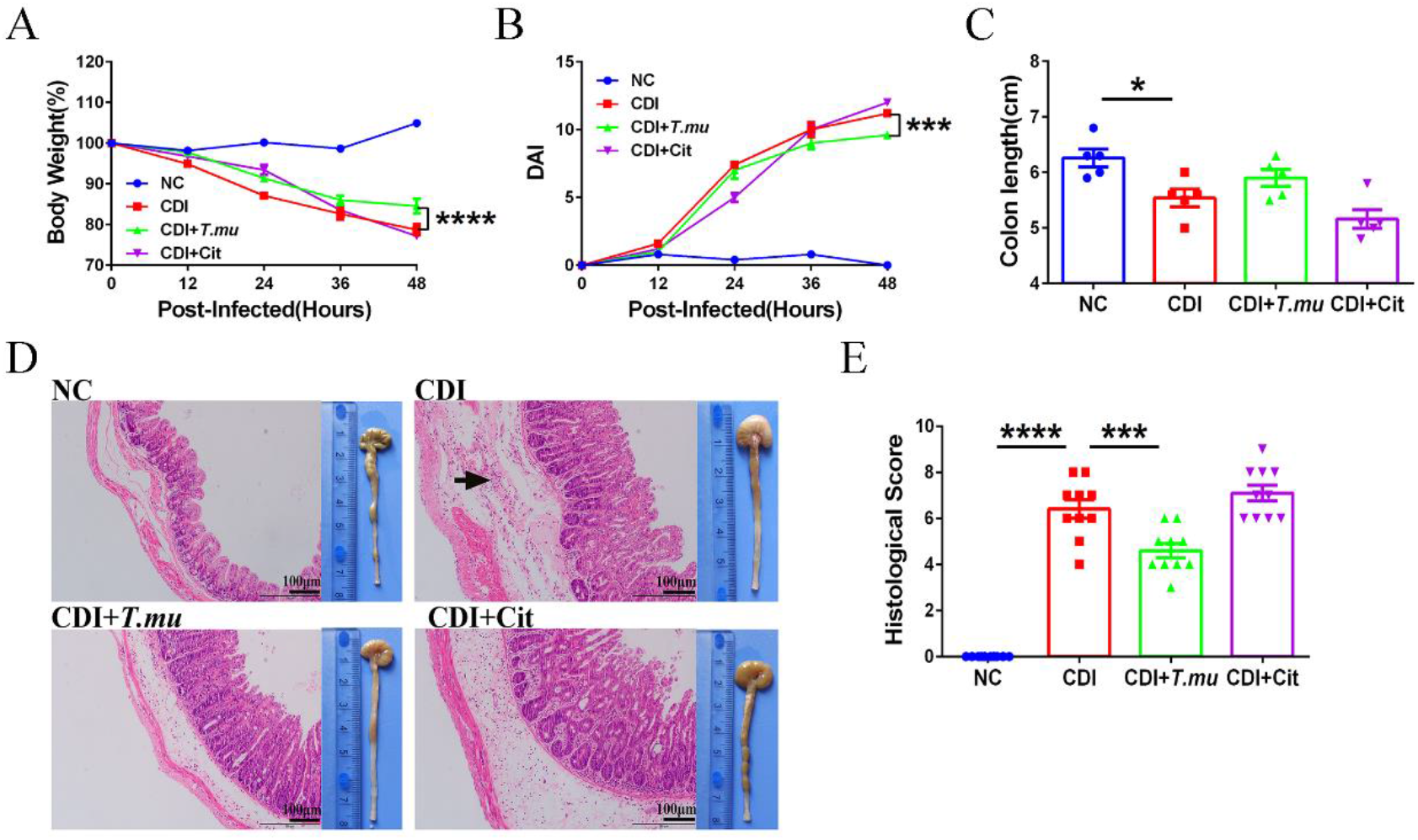
*T. mu* cannot protect intestinal damage from CDI via citrulline. (A) Body weight loss. **(B)** DAI. **(C)** Macroscopic findings of colon and representative HE-staining images (200×) of cecum. Scale bar: 100 μm. Arrow indicates infiltration of inflammatory cells. **(D)** Measurement and quantification of colon length. **(E)** Quantitation of histological score in colon. Data are the mean ± SEM. n = 5-10. Statistical significance was determined by two-way ANOVA (**A** and **B**) or one-way ANOVA (**C** and **E**). *\**: *p<*0.05. *\*\*\**: *p<*0.001. *\*\*\*\**: *p<*0.0001.

## Notes

### Competing Interest Statement

The authors have declared no competing interest.

